# Quantum chemical properties of chlorinated polycyclic aromatic hydrocarbons for delta machine learning

**DOI:** 10.1101/2024.12.19.628874

**Authors:** Dmitry Frolov, Ilya Ibraev, Igor Sedov

## Abstract

Promising Δ-machine learning approaches aim to correct the values of molecular properties obtained with computationally inexpensive methods to the accuracy of higher levels of theory. Training such models requires datasets containing the results of calculations at several different levels of quantum chemical theory. While several large and chemically diverse datasets have been published, studies in many areas require specialized datasets of structurally related molecules. Chlorinated polycyclic aromatic hydrocarbons (Cl-PAHs), the products of incomplete combustion of organic substances and materials, are hazardous pollutants with carcinogenic and mutagenic activity. Quantum chemistry methods are important to understand their formation mechanisms and properties. We describe a dataset, PACHQA, containing the results of quantum chemical calculations including properties, geometries, wavefunctions, and electron densities for 3551 molecules including 3417 Cl-PAHs with up to 6 rings and a different number of chlorine atoms in their structure as well as 134 parent polycyclic aromatic hydrocarbons (PAHs). The major part of these molecules have previously not been included in any quantum chemical datasets. The calculations were performed at three different levels of theory including geometry optimization with GFN2-xTB and r^2^SCAN-3c methods and single-point energy calculation at ωB97X-D4/def2-TZVP DFT level. The dataset can be useful to develop and validate the computational, machine learning, or experimental, approaches and study the structure-property relationships for Cl-PAHs.

## Introduction

Fast and accurate calculation of the properties of molecules is a major goal of computational chemistry. Quantum chemistry methods face a tradeoff between the speed and precision, often with a small influence of the most computationally demanding terms on the obtained geometries, energies, or other properties. The idea of replacing the most time-consuming parts of quantum chemical calculations with machine learning (ML) methods was explored in a number of studies in the past decade. It can be implemented in very different ways: for example, using ML instead of solving the Kohn-Sham DFT equations^1–3^, to describe the post-Hartree−Fock^4–6^ or DFT^7^ correlation energy using features based on the Hartree−Fock molecular orbitals, or to correct the values of properties obtained with computationally inexpensive methods to the accuracy of a higher level of theory. The latter approach was named Δ-machine learning (Δ-ML)^8^. At the same time, ML models describing large datasets of the results of quantum chemical calculations without using any input information from quantum chemistry have been actively developed^9–18^.

The term Δ-ML and the approach itself became rather popular. It has been used to improve the energetic and thermochemical properties obtained using semi-empirical methods^8,19^ or Density-Functional-Tight-Binding (DFTB) theory^20^ to DFT level, learn the difference between CCSD(T) and DFT-based potential energy surfaces^21^ or between quantum mechanical and classical potentials and forces for molecular dynamics simulations^22,23^, refine the values of redox potentials and UV-Vis absorption energies^24^, nuclear magnetic resonance chemical shifts^25,26^, dielectric properties and Raman Spectra^27^, activation energies of chemical reactions^28,29^. The ML approaches applied for Δ-ML vary from linear and kernel ridge regressions to sophisticated neural networks. They may use a variety of molecular/atomic representations and input features. A generalization of the Δ-ML concept is multifidelity machine learning^30,31^, which is based on the data from a ladder of calculation methods with increasing accuracies (called fidelities) and decreasing number of the training data points obtained using a corresponding method. Alternatively, the results of calculations at two or more lower theory levels can be used in multitask ML in which secondary regression tasks between the data at these levels provide additional information for the primary regression task^32^. Unlike most of the multifidelity approaches, non-nested training data can be used for multitask learning, i.e. the training set containing a molecule with known higher accuracy data does not necessarily include the data for this molecule obtained at all lower levels of theory. This allows one to combine the results from different independent datasets in an efficient way.

A number of large datasets of organic molecules containing results of quantum chemical calculations at two different levels of theory has been published including PubChemQC B3LYP/6-31G*//PM6^33^, QMugs^34^, tmQM^35^. Original papers include some comparisons between the values of the properties obtained using fast semi-empirical and DFT methods. Further development of Δ-ML models using these datasets is also possible. Preparation of big datasets containing structurally diverse molecules is hampered by the computational complexity of high-precision quantum chemistry methods. Moreover, the models trained on them are likely to perform badly for the molecules from the poorly represented parts of chemical space. At the same time, there are the numerous problems linked with predicting the properties for the huge series of the structurally relevant compounds. In such cases, more specialized Δ-ML models must be trained using a relevant dataset.

Chlorinated polycyclic aromatic hydrocarbons (Cl-PAHs) are the products of incomplete combustion of organic substances and materials such as polyvinyl chloride in solid wastes or dichloroethane in automotive fuels^36,37^. Cl-PAHs can also be found in chlorinated tap water. They are hazardous pollutants with carcinogenic and mutagenic properties^38,39^. Cl-PAHs are always present in the environment together with polycyclic aromatic hydrocarbons (PAHs), which are toxic and carcinogenic themselves and can transform into more toxic Cl-PAHs through chlorination^40–42^.

Accurate quantum chemistry methods are necessary to understand the mechanisms of formation and reactions of Cl-PAHs. In particular, they can provide the values of the standard thermodynamic functions of formation in the gas phase. These values can be useful for chemical kinetic models aiming to describe the mechanism of combustion and predict the concentrations of different Cl-PAHs in its products. However, the chemical space of Cl-PAHs as well as PAHs that may form in combustion processes is very large, which drives an interest in the quick, but precise quantum chemical calculations. This creates an opportunity for the application of Δ-learning methods.

Recently published COMPAS dataset^43^ contains the results of quantum chemical calculations for all possible molecules of cata-condensed PAHs composed of up to 11 benzene rings. The geometry optimization and property calculations were performed using semi-empirical GFN2-xTB method for all 34072 molecules as well as their anionic and cationic forms, and at the B3LYP-D3BJ/def2-SVP level of DFT for 8678 molecules and their ions with up to 10 benzene rings. Authors have shown that linear correlations may be sufficient to correct the energetic properties such as single point energies, HOMO/LUMO energies, or ionization potentials from xTB to the B3LYP level. Subsequently, they have performed similar analysis for cata-condensed hetero-polycyclic aromatic systems^44^ and peri-condensed PAHs^45^ and correlated DFT properties with xTB properties using linear or multi-linear regressions. In another study^46^, the enthalpies of formation of more than 700 PAHs and their derivatives were calculated and compared with available experimental data. The group additivity corrections were suggested to improve the accuracy of G3MP2B3 composite quantum chemistry method. The properties of Cl-PAHs remain less studied than those of PAHs. The work of Xu et al.^47^ comprising the results of calculations of the carbon–halogen bond dissociation energies and enthalpies of formation at 298 K for 27 monochlorinated and 27 monobrominated polycyclic aromatic hydrocarbons at different levels of theory is probably the most comprehensive study to the date.

Binding to the aryl hydrocarbon receptor (AhR), a transcription factor that regulates gene expression, is thought to be an important mechanism of toxicity of both PAHs and Cl-PAHs^48,49^, while other mechanisms are also possible^50^. Analysis of the structure–activity relationships showed that the binding affinity and toxicity correlate with the molecular properties of Cl-PAHs that can be calculated by quantum chemistry methods such as HOMO and LUMO energies, HOMO-LUMO gap, dipole moment, electronegativity, chemical hardness, electrophilicity^51–53^. This provides additional motivation for quantum chemical studies of such compounds.

The present dataset, PACHQA, is a result of quantum chemical calculations for a set of Cl-PAHs and their parent PAHs at three different levels of theory. Two of them, semi-empirical GFN2-xTB and the composite DFT r2SCAN-3c methods are used for both geometry optimization and property calculation, while the ωB97X-D4/def2-TZVP level is used for single-point calculations after r2SCAN-3c optimization. Using the dataset, it is possible to explore the Δ-learning concept as well as multitask learning approaches in order to correct various calculated properties to higher theory level.

## Methods

### Molecule selection and preparation

The chemical space of carbon skeletons in our dataset was limited to that of the rather small Cl-PAHs and PAHs included in the PubChem compounds database^54^. For the first subset (named *PubChemPCH*), 473 stable neutral molecules with at least 1 chlorine atom, 6 to 26 carbon atoms all belonging to benzenoid aromatic rings, no atoms other than C, H, and Cl, no rotatable bonds, no seven- or larger-membered rings, and no more than 6 rings in total were extracted from PubChem. The second subset, *PubChemPAH*, comprised 134 PAHs from PubChem selected using the same rules except that chlorine atoms were not allowed. This subset already included all parent PAHs for Cl-PAHs from *PubChemPCH*. Three further subsets were then constructed. The subset *monoCl* (1281 molecules) contains all possible monochlorinated derivatives of the hydrocarbons from *PubChemPAH*. The subset *perCl* (122 molecules) includes perchlorinated hydrocarbons from *PubChemPAH*, and the subset *polyCl* (1541 molecules) contains one random isomer of C_*x*_H_*y-n*_Cl_*n*_ for each hydrocarbon C_*x*_H_*y*_ from *PubChemPAH* and each integer value of *n* from 2 to *y*–1. The molecules already included in *PubChemPCH* were excluded from the *monoCl* and *perCl* subsets and were not allowed to be generated when the *polyCl* subset was constructed. The full dataset contains 3551 molecules. In Figure 1, the distribution of all the molecules in terms of the number of carbon and chlorine atoms, rings, and molecular weight is shown. The RDKit library^55^ was used to filter out or generate molecules during the dataset preparation process.

**Figure 1.**
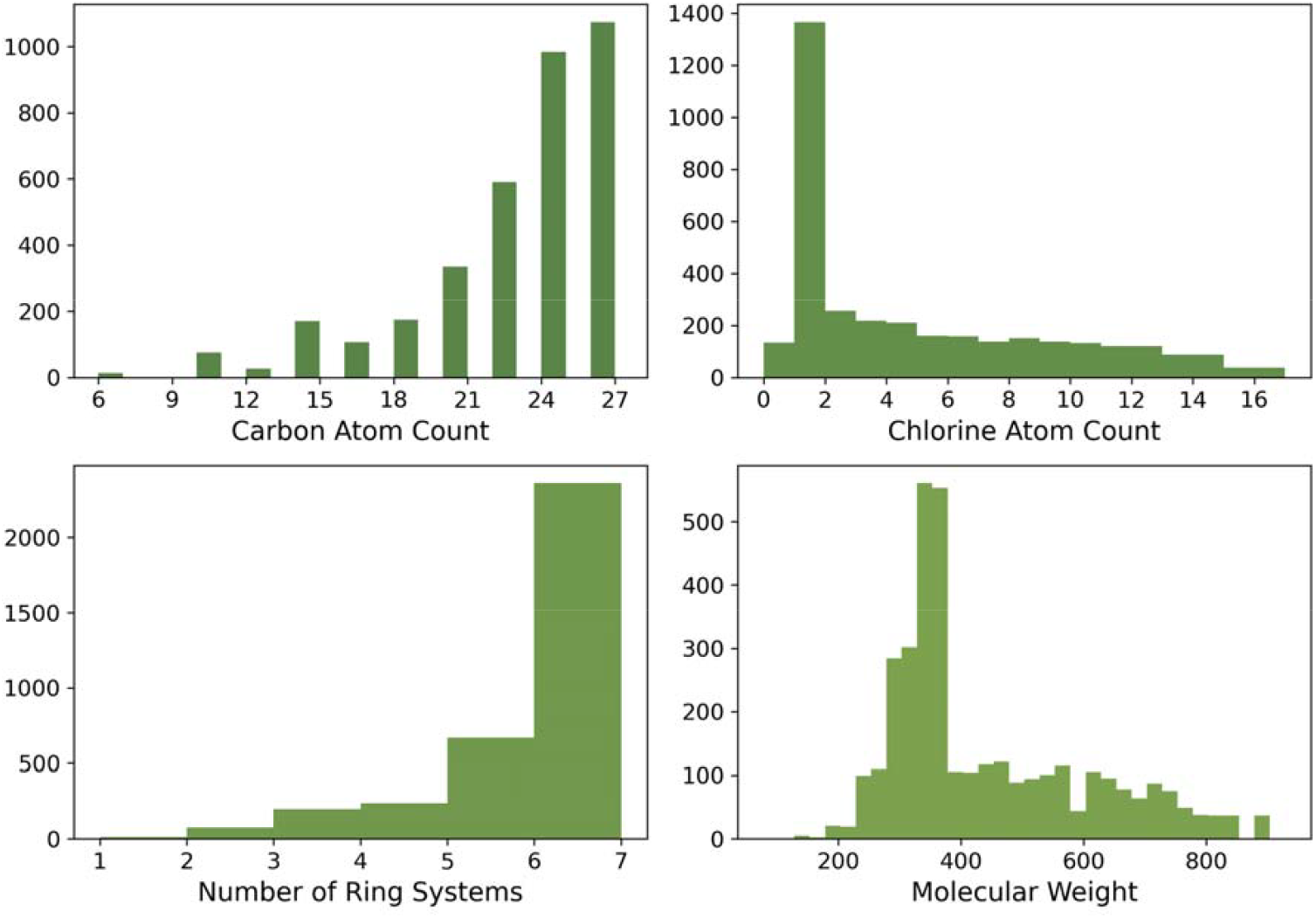
The distribution of 3551 molecules in the PACHQA dataset with respect to the number of carbon and chlorine atoms, rings, and molecular weight (g·mol^−1^).

Two structurally distinct groups of compounds can be distinguished in the dataset and all its subsets. The first group includes the compounds containing only six-membered rings. The compounds in the second group also contain at least one embedded four- or five-membered ring. Figure 2 shows the number of compounds belonging to each group in each subset as well as the examples of their molecular structures.

**Figure 2.**
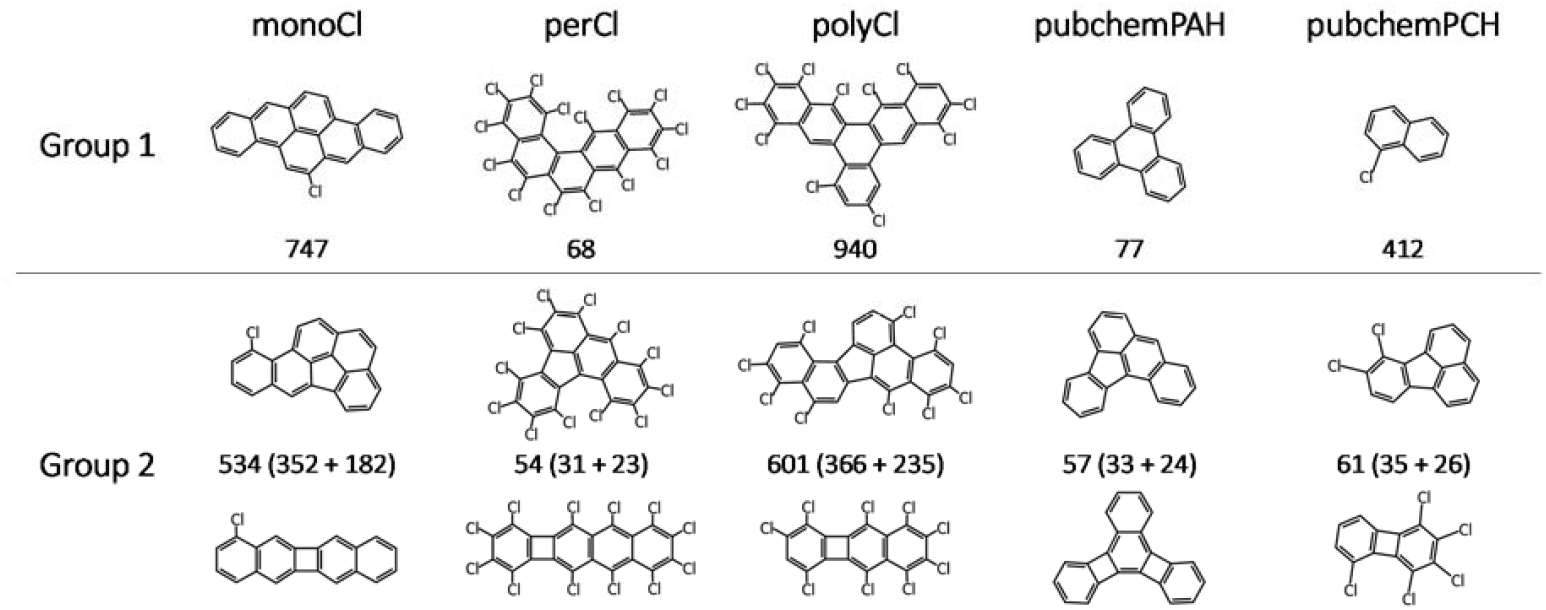
Examples of molecules from the dataset belonging to five subsets and two groups distinguished by the size of the smallest ring (Group 1 – 6, Group 2 – 5 or 4). The numbers indicate the total number of molecules in each subset and group. For Group 2, the format is “total count (5-membered smallest rings count + 4-membered smallest rings count)”.

**Figure 3.**
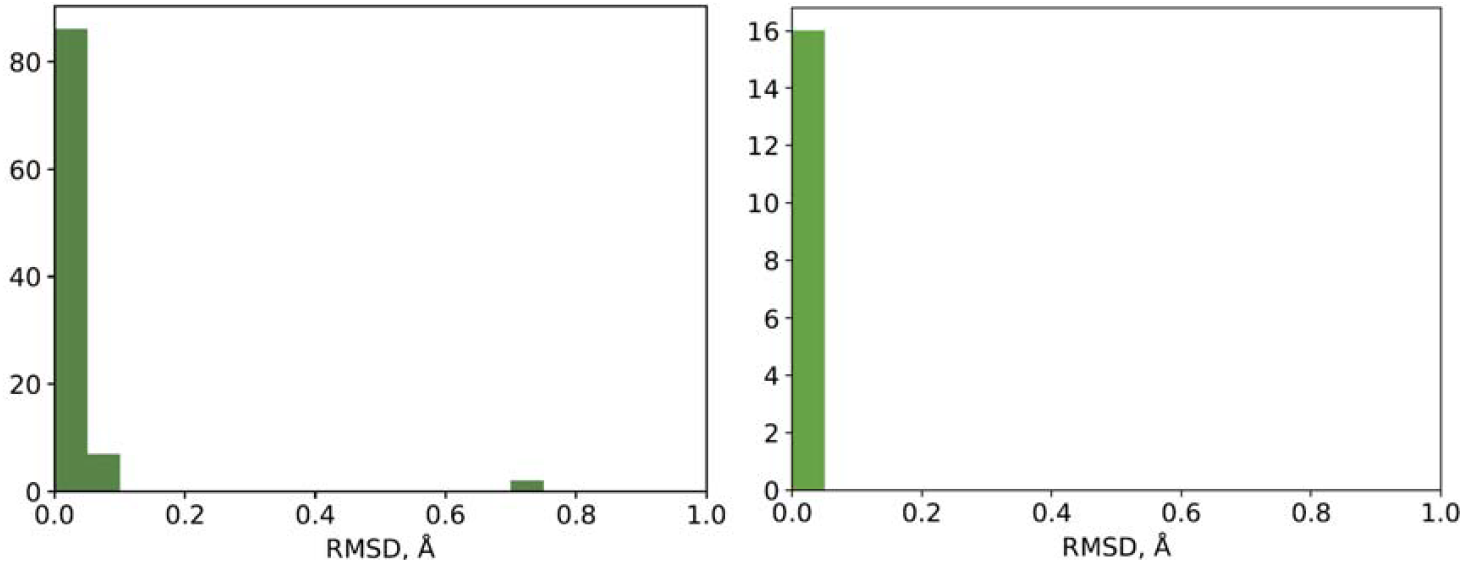
The distribution of symmetry-corrected heavy atom RMSD values between the xTB theory level structures of the same molecules in our and literature datasets. Left: molecules from the COMPAS-2x and COMPAS-3x datasets, right: molecules from the QMugs dataset.

The geometries of all molecules were preliminarily optimized in the MMFF94 force field^56^ using the steepest descent method with a maximum of 10000 steps via the OpenBabel 3.0.0 package^57^. All the molecules studied have no rotatable bonds and many of them exist as a single planar conformer. The sterically hindered molecules adopt non-planar geometry, but even in this case there are often only two energy minima corresponding to the enantiomeric conformers with the same properties. Therefore, the conformer search is not relevant for the molecules studied.

### Quantum chemical calculations

As a first step, the geometries of the molecules were optimized with the semiempirical tight-binding GFN2-xTB method^58^ using the xtb 6.6.1 package at the “extreme” optimization level with an energy convergence threshold of 5·10^−8^ Hartree. The vibrational frequencies and thermochemical properties were calculated, if imaginary frequencies appeared, the molecule was optimized again starting from the geometry distorted along the imaginary mode. The procedure was repeated until the imaginary frequencies disappeared.

The second step involved further geometry optimization using the r^2^SCAN-3c^59^ composite density functional method implemented in the ORCA 5.0.4^60^. Vibrational frequencies and thermochemical properties were also calculated, and unwanted imaginary frequencies were removed by repeated calculations starting from the geometry distorted in the corresponding direction generated by the pyQRC^61^.

In the third step, single-point energy calculations were performed for r^2^SCAN-3c geometries using the range-separated hybrid ωB97X-D4^62^ functional and the def2-TZVP basis set^63^ with the RIJCOSX approximation^64^ and the def2/J auxiliary basis set for Coulomb fitting.

### Data analysis

The Python libraries scikit-learn^65^, XGBoost^66^, PyTorch Geometric^67^, and RDKit^55^ were used for data analysis and implementation of machine learning models during technical validation.

## Data Records

The summary of molecular property values obtained after each calculation step is provided in .csv format (*props*.*csv*) in the Supplementary information. Table 1 shows the description of the data fields in this file. For each molecule, there are three records reporting the properties calculated at three different theory levels. If the property does not depend on the theory level such as the molar mass, the same value is repeated. As the Hessians have not been calculated at the ωB97X-D4/def2-TZVP level, the information on vibrational frequencies for thermochemical calculations at this level was taken from the r^2^SCAN-3c results.

**Table 1.** The properties of the molecules stored in the *props*.*csv* file.

| Property | *props.csv* field | Units |
| --- | --- | --- |
| Molecule identifier within the dataset | id |  |
| The subset to which the molecule belongs | subset |  |
| InChIKey | key |  |
| SMILES | smiles |  |
| Molecular formula | formula |  |
| The level of theory at which the properties have been calculated. GFN2-xTB, r<sup>2</sup>SCAN-3c, and ωB97X-D4/def2-TZVP are denoted as xtb2, r2scan, and d4tzvp, respectively. | level | |
| Molecular weight | mwt | $\text{g}\cdot\text{mol}^{-1}$ |
| Number of hydrogen atoms | nH |  |
| Number of carbon atoms | nC |  |
| Number of chlorine atoms | nCl |  |
| Single-point energy | E | $E_{\text{h}}$ |
| Total enthalpy at 298.15 K | H | $E_{\text{h}}$ |
| Total entropy at 298.15 K | S | $E_{\text{h}}\cdot\text{K}^{-1}$ |
| Total Gibbs free energy at 298.15 K | G | $E_{\text{h}}$ |
| Energy of atomization | dE | $E_{\text{h}}$ |
| Enthalpy of atomization at 298.15 K | dH | $E_{\text{h}}$ |
| Entropy of atomization at 298.15 K | dS | $E_{\text{h}}\cdot\text{K}^{-1}$ |
| Gibbs free energy of atomization at 298.15 K | dG | $E_{\text{h}}$ |
| Zero-point energy | zpe | $E_{\text{h}}$ |
| Energy of the highest occupied molecular orbital (HOMO) | homo | eV |
| Energy of the lowest unoccupied molecular orbital (LUMO) | lumo | eV |
| HOMO–LUMO gap | gap | eV |
| Dipole moment | Mu | D |
| Electronic polarizability | alpha | a.u. |
| The dipole moment components (*x y z*) (space delimiter) | tMu | a.u. |
| Rotational constants | rotA, rotB, rotC | $\text{cm}^{-1}$ |

The full results of the calculations are available at Science Data bank^68^. They include optimized geometries and more properties which may be useful for machine learning tasks in *PACHQA1-main*.*7z* (geometries, xtb output, ORCA property reports, 183 MB) and *PACHQA2-full_outfiles*.*7z* (full ORCA output files, 343 MB) archives.

The file *PACHQA3-wfns*.*7z* (57 GB) contains wavefunctions, electron densities, and xtb electrostatic potentials. All other files generated during calculations including the output of calculations that result in imaginary frequencies are collected in the *PACHQA4-other*.*7z* file (2 GB), which is unlikely to be demanded.

## Technical Validation

The dataset shares no or very few molecules with some large datasets of quantum chemical properties because these datasets contain only drug-like compounds (Alchemy^69^, Aquamarine^70^, SPICE^71^) or include only molecules with a small number of heavy atoms (QM7-X^72^, QM9^73^, Vector-QM24^74^). The PubChemQC database^33^ containing the results of calculations for the largest number (86 million) of different organic molecules listed on PubChem includes the majority of molecules in *PubChemPAH* and *PubChemPCH* subsets and no molecules from the other subsets. The COMPAS family datasets containing the properties of PAHs have common molecules with *PubChemPAH* subset: 58 molecules in COMPAS-1x and COMPAS-1D^43^ (57 of these molecules with the exception of benzene are also found in COMPAS-2x), 76 molecules in COMPAS-2x^44^, and 19 molecules in COMPAS-3x and COMPAS-3D^45^. The QMugs dataset^34^ shares 8 molecules with *PubChemPAH* and 8 molecules with the *PubChemPCH* subset. Both COMPAS and QMugs datasets include the results obtained using xTB method and are suitable to validate our calculations.

The geometry of common molecules in different datasets was compared by calculating the symmetry-corrected heavy atom RMSD values. Due to the possibility of existence of enantiomeric conformers of non-planar molecules with the same energy, the RMSD was calculated for both enantiomers, and the lower value was taken. Only 2 molecules out of 95 in COMPAS-2x and COMPAS-3x datasets had a significantly deviating geometry with RMSD > 0.1 Å, which is explained by the rather rare case of “diastereomeric” conformers of non-planar PAH molecules. In the Qmugs dataset, all 16 common molecules showed the RMSD below 0.05 Å.

The physical properties of these molecules calculated using the GFN2-xTB method in our and literature datasets are also in agreement. Table 2 shows the values of maximum deviations for different properties of all the molecules shared by QMugs, COMPAS-1x and COMPAS-3x datasets except for the molecules with significantly different geometry mentioned above. (The COMPAS-2x dataset uses a different GFN1-xTB method, so the properties have larger deviations from our results.)

**Table 2.** The maximum deviations of various properties calculated using the GFN2-xTB method for the molecules shared between our and literature datasets (QMugs, COMPAS).

| Property | Max deviation |
| --- | --- |
| QMugs dataset |  |
| Single-point energy | 0.11 kJ·mol<sup>-1</sup> |
| Total enthalpy at 298.15 K | 0.11 kJ·mol<sup>-1</sup> |
| Total entropy at 298.15 K | 0.53 J·K<sup>-1</sup>·mol<sup>-1</sup> |
| Total Gibbs free energy at 298.15 K | 0.14 kJ·mol<sup>-1</sup> |
| HOMO energy | 6.4·10<sup>-4</sup> eV |
| LUMO energy | 2.3·10<sup>-3</sup> eV |
| HOMO–LUMO gap | 1.7·10<sup>-3</sup> eV |
| Dipole moment | 0.023 D |
| Polarizability | 4.6·10<sup>-4</sup> a.u. |
| COMPAS-1X and COMPAS-3X datasets |  |
| Single-point energy | 0.12 kJ·mol<sup>-1</sup> |
| HOMO energy | 2.6·10<sup>-3</sup> eV |
| LUMO energy | 5.4·10<sup>-3</sup> eV |
| HOMO–LUMO gap | 8.1·10<sup>-3</sup> eV |
| Dipole moment | 4·10<sup>-3</sup> D |
| Zero-point energy | 0.07 kJ·mol<sup>-1</sup> |

The values of the electronic polarizabilities calculated at two higher levels of theory show a good agreement with the experimental data for 20 molecules from the compilation of Gussoni et al.^75^ (Figure 4). The RMSD values are 27.7 a.u. at the GFN2-xTB level, 6.7 a.u. at the r2SCAN-3c level, and 6.6 a.u. at the ωB97X-D4/def2-TZVP level.

**Figure 4.**
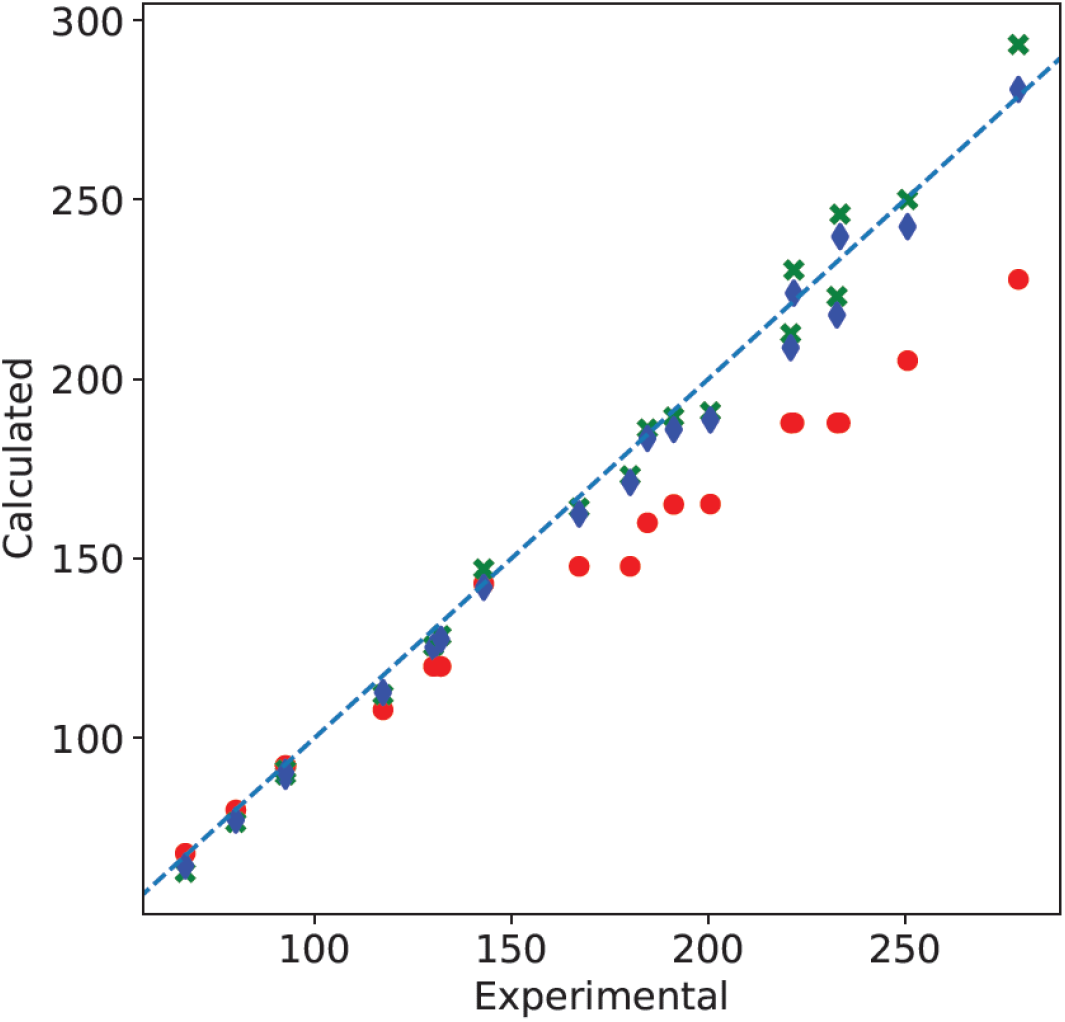
Calculated (red circles – GFN2-xTB, green crosses – r2SCAN-3c, blue triangles – ωB97X-D4/def2-TZVP) vs experimental^75^ values of electronic polarizabilities (a.u.). The line corresponds to the *y* = *x* equation.

The statistics of the changes in molecular geometries during the semi-empirical and composite DFT optimizations is visualized in Figure 5 as the distribution of the symmetry-corrected heavy atom RMSD values. The GFN2-xTB optimization resulted in the larger changes of the initial structures corresponding to the energy minima in the MMFF94 force field than the subsequent r^2^SCAN-3c step which resulted mainly in fine geometry adjustments. The connectivity of all the molecules was preserved during the geometry optimizations and is identical in the original .sdf files and in the molecular graphs generated by RDKit from the atomic coordinates after the final r^2^SCAN-3c optimization.

**Figure 5.**
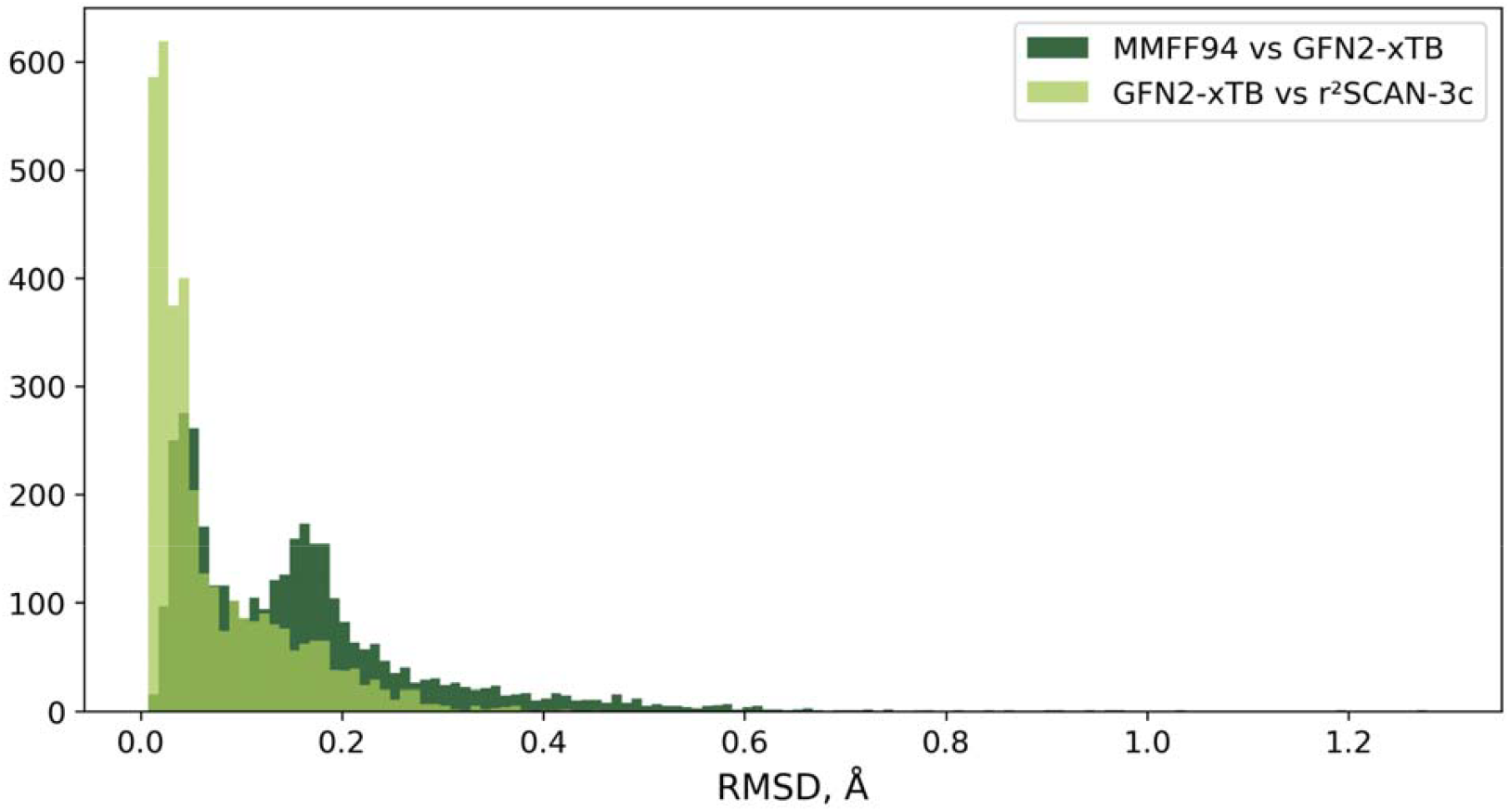
The distribution of symmetry-corrected heavy atom RMSD values between the structures after different steps of geometry optimization.

Changes in atomic positions during geometry optimization are often related to changes in molecular planarity. The average deviation of coordinates of heavy atoms from the plane of best fit (PBF)^76^ is a simple measure of planarity. The distribution of PBF scores after the geometry optimization at each level of theory is shown in Figure 6a. The most obvious steric clashes become resolved after MMFF94 minimization, but a significant number of molecules lose their planarity at GFN2-xTB or even r^2^SCAN-3c step. Approximately one third of all molecules in the dataset can be described as planar after the r^2^SCAN-3c optimization, while others show more or less significant deviations from planarity in their energy minimum. The majority of hydrocarbon molecules from *PubChemPAH* subset remain planar (Figure 6b), while the introduction of a single chlorine atom significantly increases the steric hindrance, leading to a larger fraction of non-planar molecules in the *monoCl* subset. Consequently, the vast majority of perchlorinated compounds in *perCl* subset are far from planar. This can be also demonstrated by the principal moments of inertia plots (Figures 3c, 3d).

**Figure 6.**
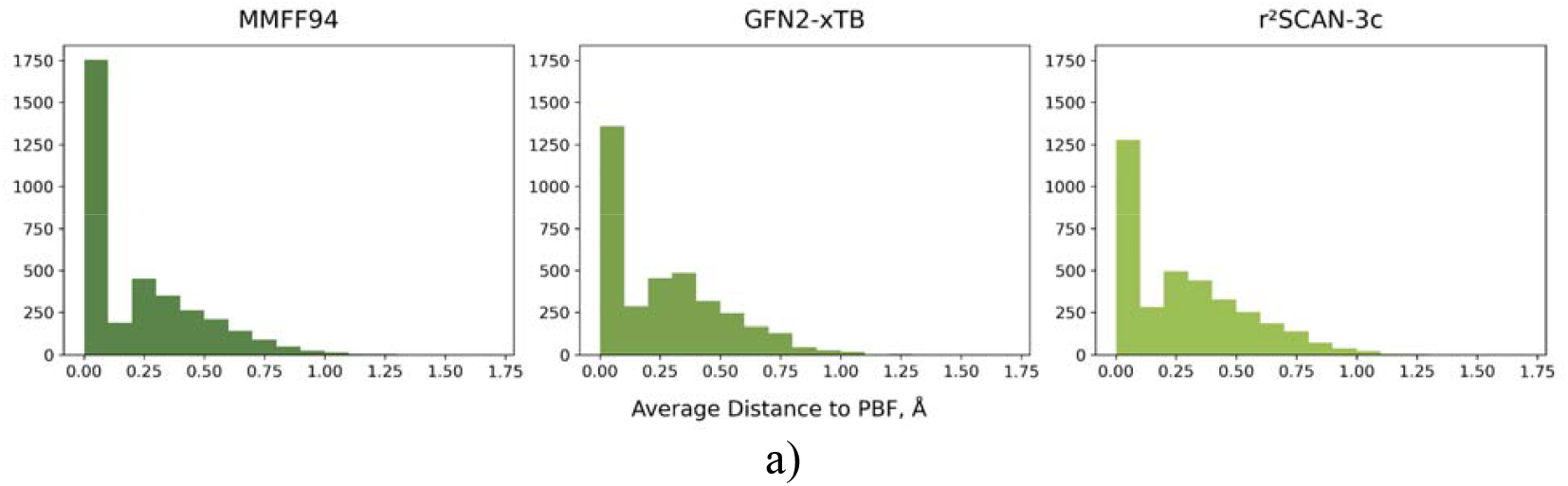
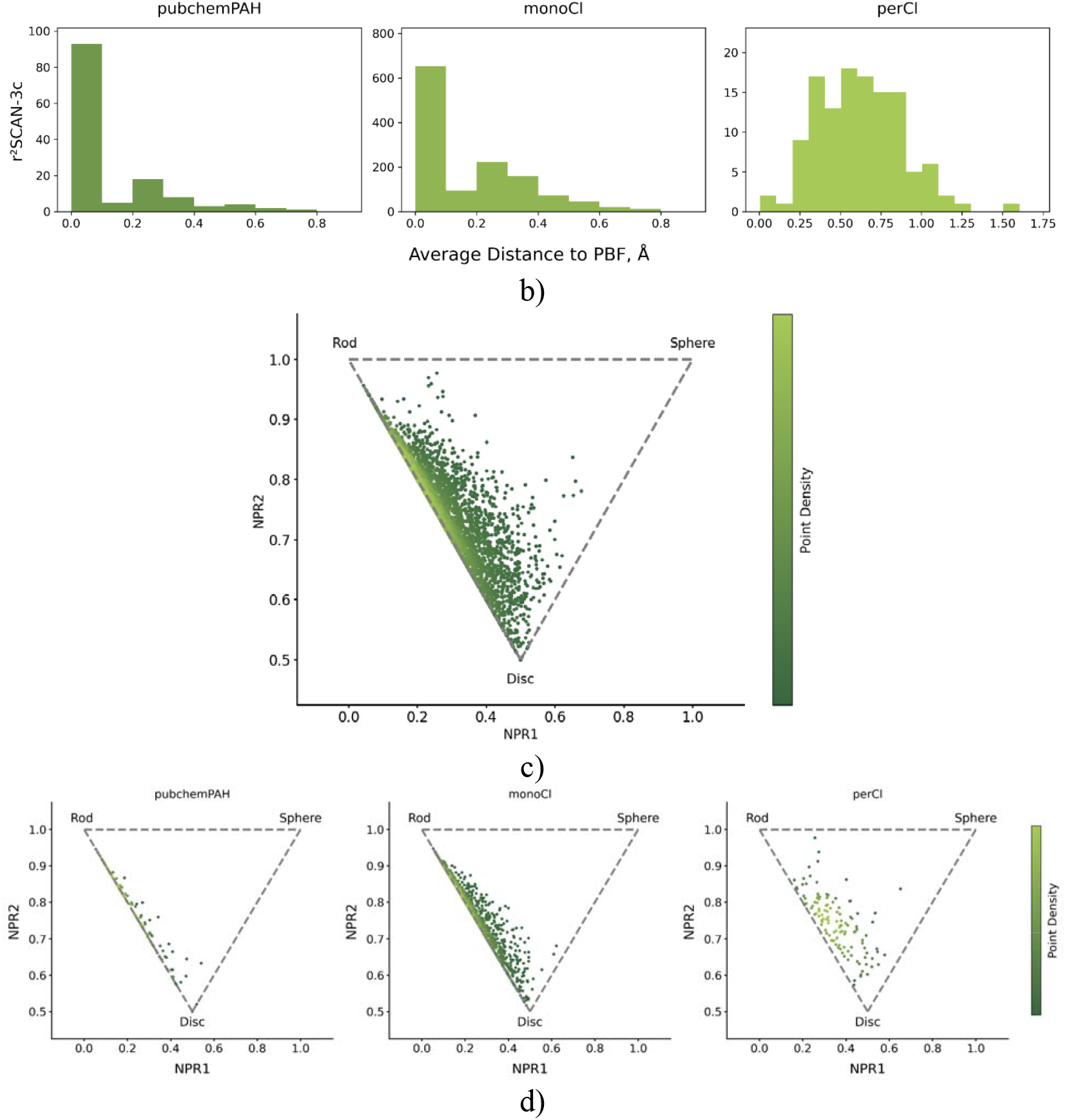
a) The distribution of plane of best fit (PBF) scores after the geometry optimization at each level of theory. b) The distribution of PBF scores in *PubChemPAH, monoCl*, and *perCl* subsets after r^2^SCAN-3c geometry optimization. c, d) The principal moments of inertia plot for a complete dataset (c) and its subsets (d).

The total time of GFN2-xTB geometry optimization and frequency calculation for a complete dataset was 58 core hours, which is 220 times less than the total time of r^2^SCAN-3c optimization and frequency calculation (12761 core hours, this value does not include the time required to get rid of imaginary frequencies, which additionally took about 6% of this time). Single-point energy calculations at the ωB97X-D4/def2-TZVP level required 3816 core hours. Thus, the total time of the DFT calculations was 16577 core hours, 286 times more than at the semi-empirical level. Figure 7 shows the relationships between the computational costs at different theory levels for individual molecules.

**Figure 7.**
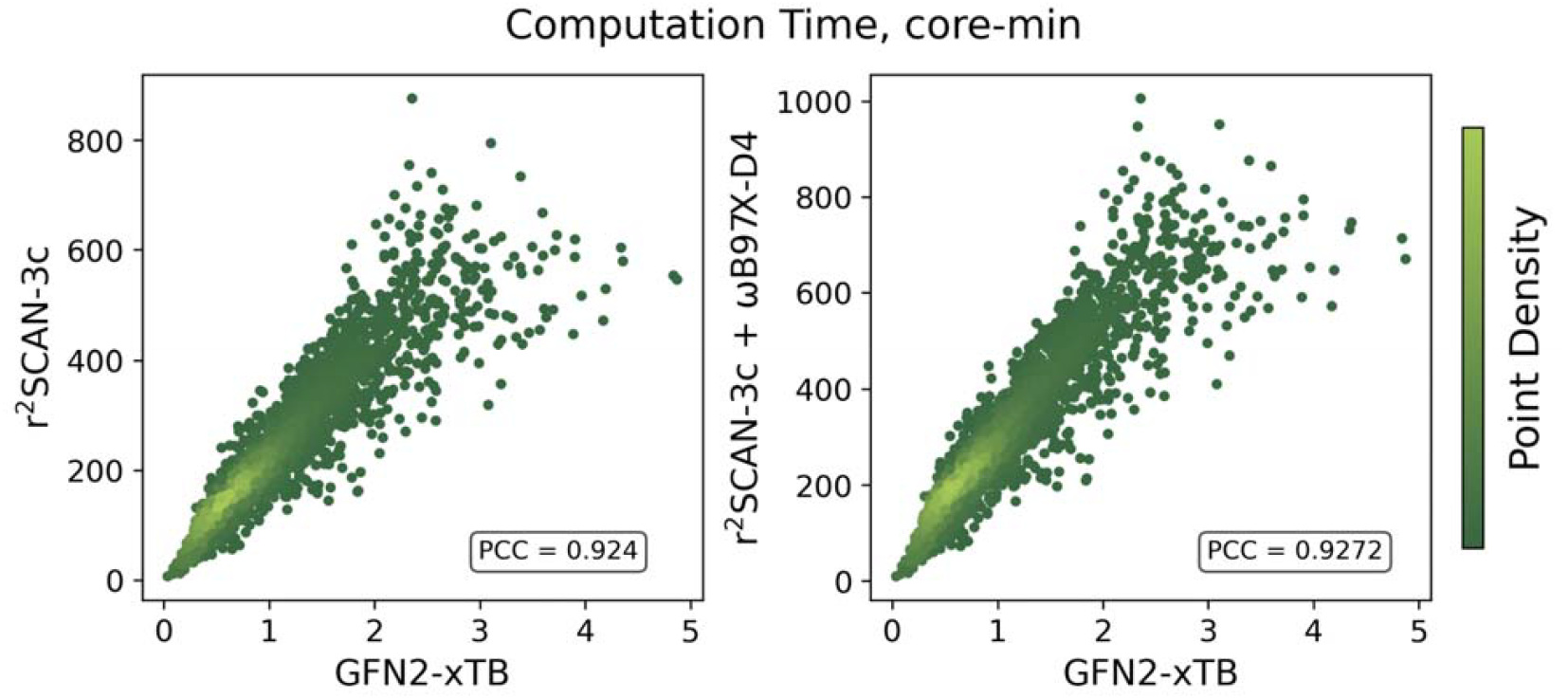
Comparison of the computational costs at different theory levels for individual molecules: a) r^2^SCAN-3c vs GFN2-xTB optimization and frequency calculation; b) r^2^SCAN-3c optimization and frequency calculation with subsequent ωB97X-D4/def2-TZVP single-point energy calculation vs GFN2-xTB optimization and frequency calculation.

The use of Δ-ML instead of high theory levels has a potential to become a key for fast and accurate computations. Figure 8 compares the values of some properties obtained using different quantum chemical methods. Even the simplest model, a linear regression between the values at different levels of theory, gives a satisfactory description of some higher level properties such as the HOMO–LUMO gap. More sophisticated models will be required for accurate prediction of enthalpy of atomization or electronic polarizability. The parameters of the linear fits for the properties shown in Figure 8 are summarized in Table 3.

**Table 3.** The parameters of the linear regressions *y* = *ax* + *b* between the properties calculated at different levels of theory for a complete dataset (3551 points)

| Property | $a$ | $b$ | RMSD | $R^2$ |
| --- | --- | --- | --- | --- |
| r<sup>2</sup>SCAN-3c vs GFN2-xTB |  |  |  |  |
| Enthalpy of atomization | 0.78 | −0.0164 | 0.0384 $E_h$<br>(101 kJ·mol<sup>−1</sup>) | 0.9991 |
| HOMO–LUMO gap | 1.16 | −0.005 | 0.083 eV | 0.9751 |
| Dipole moment | 0.92 | 0.028 | 0.153 D | 0.9757 |
| Polarizability | 1.42 | −33.3 | 19.9 a.u. | 0.9618 |
| ωB97X-D4/def2-TZVP vs r<sup>2</sup>SCAN-3c | | | | |
| Enthalpy of atomization | 0.98 | 0.0242 | 0.0211 $E_h$<br>(55 kJ·mol<sup>−1</sup>) | 0.9997 |
| HOMO–LUMO gap | 1.32 | 4.09 | 0.127 eV | 0.9673 |
| Dipole moment | 1.03 | 0.012 | 0.102 D | 0.9899 |
| Polarizability | 0.89 | 20.7 | 4.0 a.u. | 0.9981 |
| ωB97X-D4/def2-TZVP vs GFN2-xTB | | | | |
| Enthalpy of atomization | 0.77 | 0.0048 | 0.0213 $E_h$<br>(56 kJ·mol<sup>−1</sup>) | 0.9997 |
| HOMO–LUMO gap | 1.53 | 4.08 | 0.160 eV | 0.9481 |
| Dipole moment | 0.95 | 0.046 | 0.202 D | 0.9602 |
| Polarizability | 1.26 | −9.7 | 16.9 a.u. | 0.9652 |

**Figure 8.**
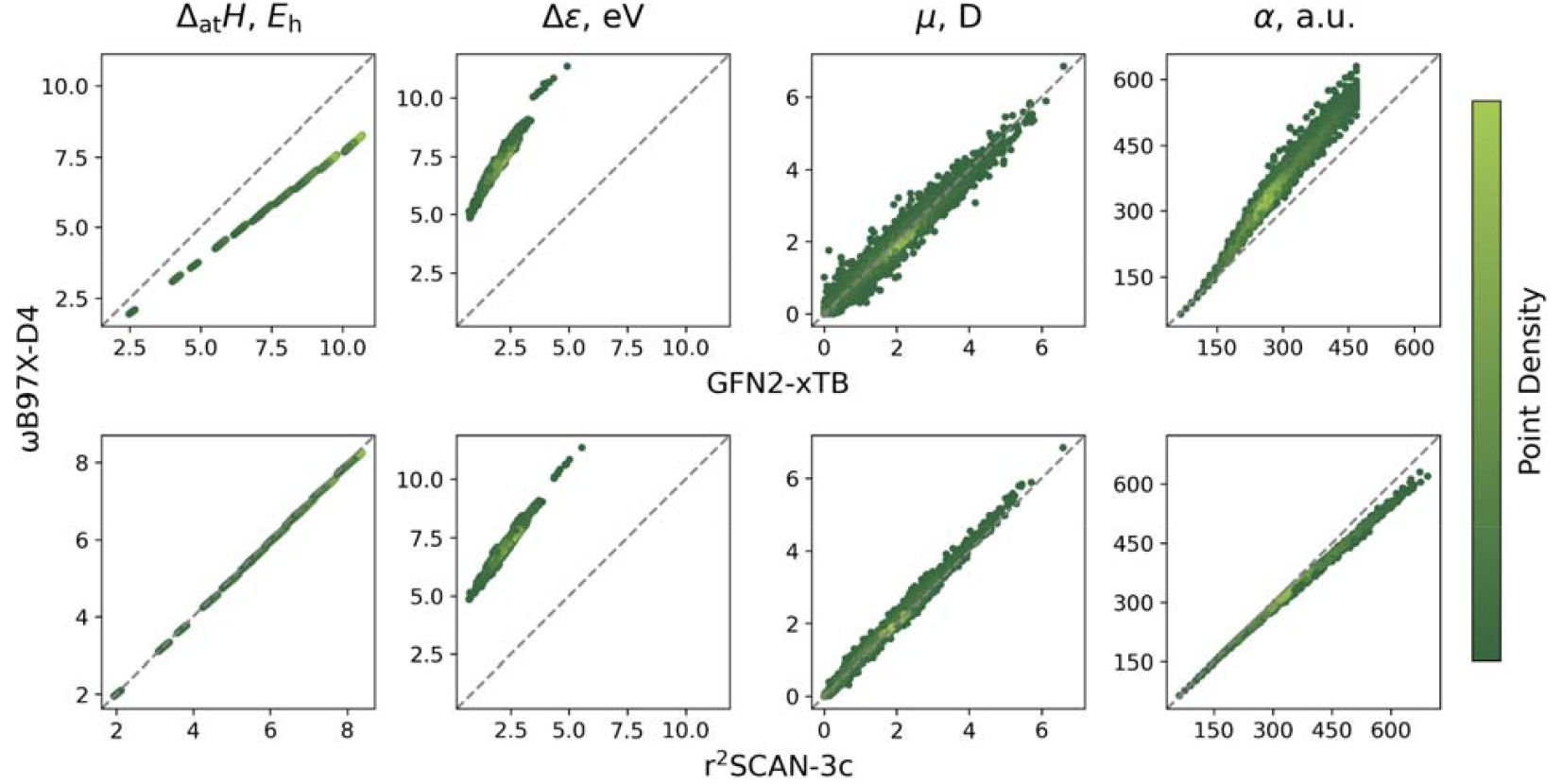
Comparison of the values of enthalpy of atomization at 298.15 K (Δ_at_*H*), HOMO–LUMO gap (Δε), dipole moment (μ), and electronic polarizability (α) obtained at higher (ωB97X-D4/def2-TZVP) and lower (GFN2-xTB, r^2^SCAN-3c) levels of theory.

## Supporting information

Summary of calculated molecular property values

## Acknowledgements

This work was supported by the grant of the state program of the «Sirius» Federal Territory «Scientific and technological development of the «Sirius» Federal Territory» (Agreement №18-03 date 10.09.2024).

## Author Contributions

Dmitry Frolov: Investigation, Software, Formal Analysis, Data Curation, Visualization, Writing – Original Draft.

Ilya Ibraev: Investigation, Visualization.

Igor Sedov: Conceptualization, Methodology, Investigation, Software, Formal Analysis, Writing – Original Draft, Supervision.

